# An infection assay recapitulating *Xanthomonas translucens* pv. *translucens* natural entry into barley hydathodes

**DOI:** 10.64898/2026.10.06.757142

**Authors:** Alain Jauneau, Aude Cerutti, Jules Butchacas, Marie-Christine Auriac, Jean-Marc Routaboul, Claude Bragard, Jonathan M. Jacobs, Laurent D. Noël

## Abstract

Pathogenic bacteria navigate biological host landscapes to enter and establish infection. How *Xanthomonas* phytopathogenic bacteria enter specific plant tissues is believed to directly determine the systemic vs. local outcome of infection. Non-vascular *Xanthomonas* infect leaf mesophyll locally from stomatal pores, while vascular ones enter through hydathode water pores on the leaf margin before invading the xylem for systemic movement. The pathovar *Xanthomonas translucens* pv. *translucens* (*Xtt*) is associated with barley infection, causing both bacterial leaf streak and bacterial blight symptoms by colonizing both the mesophyll spaces through stomatal guard cells and the plant vasculature, respectively. How *Xtt* reaches the vasculature in barley leaves is yet to be established. In this study, we determined the anatomy of barley hydathodes. A protocol allowing efficient hydathode infection by *Xtt* was established and caused bacterial blight symptoms, closely mimicking the natural infection process. This study establishes *Xtt* as one of the few demonstrated hydathode pathogens.

## Introduction

Plant pathogens in the genus *Xanthomonas* are bacteria that cause diseases in over 400 plant species (Jacques et al. 2016; Timilsina et al. 2020). These bacteria typically enter their hosts through natural openings such as pores or wounds and exhibit either vascular or non-vascular tissue-specific colonization behaviours. Non-vascular *Xanthomonas* typically enter the host via stomata, which are found on leaf surfaces and play a critical role in gas exchange (e.g. CO2, O2, H2O, Clark et al. 2022; Edwards et al. 1998). Vascular *Xanthomonas* pathogens move long distances within the host xylem and cause widespread, systemic, and seed-borne infections causing complete leaf blight, wilting, and even death of the plant (Yadeta and Thomma 2013). Significant vascular diseases caused by *Xanthomonas* include, but are not limited to, bacterial blight (BB) of rice caused by *Xanthomonas oryzae* pv. *oryzae* (*Xoo*) (Nino-Liu et al. 2006), black rot of cabbage caused by *Xanthomonas campestris* pv. *campestris* (*Xcc*) (Smith 1897) and BB of barley caused by *Xanthomonas translucens* pv. *translucens* (*Xtt*) (Jones et al. 1916). While the former two are recognized for accessing the host vasculature through hydathode organs (Guo and Leach 1989; Hugouvieux et al. 1998; Tabei and Mukoo 1960), the way *Xtt* gets access to plant vasculature has not been reported.

Hydathodes are organs present on the leaves of vascular plants and play a critical role in guttation, a process involving the release of xylem-derived fluids during periods of high humidity and low transpiration (Cerutti et al. 2019; Haberlandt 1897; Singh 2013). Although they have long remained poorly characterized, recent studies have provided new insights into their anatomy, development and physiology (Bellenot et al. 2022; Cerutti et al. 2017; Dieffenbach et al. 1980; Jauneau et al. 2020; Nagai et al. 2013; Perrin 1972; Routaboul et al. 2024; Savourat et al. 2026). Epithemal hydathodes (formerly called passive hydathodes) typically consist of terminal tracheids, epithem cells, and an epidermis with large water pores that resemble stomata (e.g., Cerutti et al. 2017; Jauneau et al. 2020; Perrin 1972).

Because hydathodes are tightly connected to xylem vessels, they serve as the primary entry point for several vascular phytopathogenic bacteria. In the xylem, these pathogens can spread and cause systemic infections in plants. Examples include *Xanthomonas axonopodis* pv. *dieffenbachiae* (Anthurium, Toves 2008), *Clavibacter nebraskensis* (maize, Mullens and Jamann 2021), *Clavibacter michiganensis* subsp. *michiganensis*, (tomato, Carlton et al. 1998), *Xanthomonas hortorum* pv. *gardneri* (tomato, Bernal et al. 2021), *Xoo* (Guo and Leach 1989; Tabei and Mukoo 1960) and *Xcc* (Hugouvieux et al. 1998). *Xcc* has emerged as a model for studying hydathode infection by such vascular bacterial pathogens (Cerutti et al. 2017; Hugouvieux et al. 1998). During infection of cabbage and Arabidopsis by *Xcc*, hydathodes deploy active immune responses (Cerutti et al. 2017; Lauber et al. 2024; Paauw et al. 2023; Pfeilmeier et al. 2025) while bacteria produce a large number of virulence proteins including type III effector proteins (Cerutti et al. 2017) and plant cell wall-degrading enzymes secreted by *Xanthomonas* (Gluck-Thaler et al. 2020; Luneau et al. 2022b; Luneau et al. 2022a; Paauw et al. 2024; Rajeshwari et al. 2005; Tayi et al. 2018) which altogether determines the outcome of the interaction.

*Xtt* is a global bacterial pathogen of barley capable of causing several disease symptoms such as bacterial blight (BB), black chaff, and bacterial leaf streak (BLS). BLS causes short, water-soaked lesions on host plant leaves upon infection at stomata that later spreads to the mesophyll tissue (Shekhawat and Patel 1978). These lesions can vary in size and shape and are often surrounded by a yellow halo. Bacterial blight symptoms caused by *Xtt* are hypothetized to result from pathogen’s entry at hydathodes and subsequent colonization of the plant’s vascular system.

In monocot species such as rice, commonly used inoculation methods for vascular pathogens rely on leaf-wounding techniques such as leaf clipping, which bypass hydathodes (Gluck-Thaler et al. 2020; Kauffman et al. 1973). In this study, we describe a novel inoculation method of *Xtt* on barley that recapitulates BB symptoms from a hypothesized hydathode natural infection (Jones et al. 1916). We provide a detailed cytological description of barley hydathodes and of their infection by *Xtt*. This study establishes *Xtt* as one of the few demonstrated hydathode pathogens and elucidates how *Xtt* gets access to xylem vessels.

## Materials and Methods

### Bacterial strains, plasmids, and growth conditions

The GFP-tagged *Xtt* strain UPB886 pNEO-GFP (Gluck-Thaler et al. 2020; Han et al. 2008) used throughout this study is a derivative of UPB886 originally isolated on *Hordeum maritimum* (1990, Iran) (Bragard et al. 1997; Bragard et al. 1995). Bacteria were grown at 28°C in Nutrient Broth liquid medium (15 g/L peptone, 3 g/L yeast extract, 6 g/L NaCl, 1 g/L D(+) glucose) supplemented with kanamycin (50 µg.mL^-1^).

### Plant material, growth conditions and infection tests

Barley (*Hordeum vulgare* L cv. Morex) was grown in a growth chamber under short-day conditions (8 h light, 22°C, 70% relative humidity). Bacteria were harvested from overnight cultures by centrifugation (4,000g, 10 min) and resuspended at 10^8^ cfu.mL^-1^ in 1 mM MgCl_2_ and 0.02% SILWET L-77. Inoculations were performed on 8- to 10-day-old barley seedlings by dipping the top half of the first true leaves for 20 s in the bacterial suspension with up-and-down shaking. Plants were placed for 24 h in miniature greenhouses with a clear plastic cover (100% relative humidity) before being returned to ambient humidity.

### Clarification of leaf samples

Barley leaves were treated for several weeks in a chloral hydrate solution and mounted between a glass slide and a coverslip in the same solution as described (Cerutti et al. 2017). Nomarski differential interference contrast (DIC) images were acquired using an inverted microscope (DMIRBE, Leica, Germany) with a long-working-distance objective lens 20X NA= 0.8.

### *In vivo* confocal microscopy

Fresh leaf samples were mounted in water on a glass slide and covered with a coverslip. Confocal images were acquired with a laser scanning confocal microscope (SP2 AOBS, Leica) equipped with 40X water- or oil-immersion objective lenses as described (Cerutti et al. 2017) and a spectral module. For fluorescence imaging, calcofluor or phenolic compounds were excited using a 405-nm diode laser, and emission was collected between 410 and 500 nm. Chlorophyll was excited using the 633-nm line of a helium-neon laser, and emission was captured between 640 and 680 nm. GFP was excited via the 488-nm argon laser line, and emission was collected from 500 to 540 nm.

### Preparation of fixed embedded material for optical microscopy

Healthy barley leaf sections were fixed, dehydrated, and embedded in resin (London resin white) as described (Cerutti et al. 2017). Sections were stained with calcofluor (0.01% in aqueous solution) and rapidly rinsed. Images were acquired with a confocal microscope (SP2 AOBS, Leica). Calcofluor and phenolics were excited using a 405 nm diode laser, and emission was collected from 410 to 500 nm. Observations were made using a 40X N.A. 1.3 oil immersion lens. For better visualization, we encoded the images in grayscale and inverted colors.

Inoculated barley leaf sections were embedded in LR White resin and cut with an Ultracut E ultramicrotome equipped with a diamond knife (Reichert-Leica). Sections were stained with toluidine blue, methylene blue, and fuchsine. Images were acquired either with an inverted microscope (DMIRBE, Leica) equipped with a long-working-distance dry objective lens 40X, with a confocal microscope (Leica SP2 AOBS, Germany) using brightfield imaging and a 40X N.A. 1.3 oil immersion lens or with a Nanozoomer-HT (Hamamatsu®). We used the Image Pro-plus software to adjust contrast and refine the outlines of small objects using a Gaussian filter.

### Preparation of material for cryo-scanning electron microscopy

Fresh 10-day-old barley leaf tips were prepared and imaged by cryo-scanning electron microscopy as described (Cerutti et al. 2017).

## Results

### Recapitulation of barley hydathode infection by *Xtt* using dip inoculation

We tested a dip-inoculation method tailored to allow hydathode infection, similar to that used for hydathode infection in cauliflower or Arabidopsis by *Xcc* (Cerutti et al. 2017). Eight- to 10-day-old barley leaves (*Hordeum vulgare* cv. Morex) were dipped in a bacterial suspension of *Xtt* strain UPB886 pNEO-GFP. Two weeks after inoculation, macroscopic disease symptoms were visible (Figure 1). In contrast to the mock control, samples inoculated with *Xtt* exhibited water-soaked lesions at the leaf tip and elongated water-soaked areas along the leaf margin, reminiscent of early symptoms commonly associated with bacterial blight (BB). Under this inoculation procedure, *Xtt* preferentially causes symptoms at the leaf tip where leaf apical hydathodes are found.

**Figure 1.**
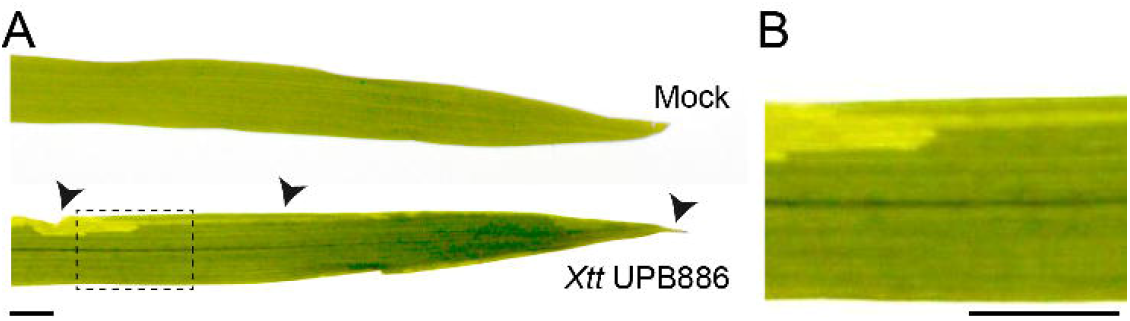
Dip inoculation protocol with *Xtt* recapitulates bacterial blight (BB) symptoms on natural infection of barley leaves. **A**, Composite image showing disease symptoms 14 days post-dip inoculation of barley leaves with mock control (1 mM MgCl_2_) and *Xtt* strain UPB886 pNEO-GFP. Arrowheads indicate water-soaked, chlorotic, and necrotic symptoms at apical hydathodes or along leaf margins. **B**, Close-up images shown in (A) for leaves inoculated with *Xtt* UPB886 pNEO-GFP (B). Scale bars: 0.5 cm.

### the anatomy of barley hydathodes is typical of epithemal hydathodes

To determine the nature of *Xtt* hydathode infection, we first describe barley hydathode anatomy. On young barley seedlings, guttation was observed only at the leaf apex of the first leaves (Figure 2A), implying that a hydathode exists at the leaf tip. Further, droplet release on both the adaxial and abaxial sides of the leaf suggests that water pores are present on both surfaces, as previously reported (Dieffenbach et al. 1980). Using cryo-prepared leaf tips and scanning electron microscopy (SEM) (Figure 2B-E), we observed numerous water pores, particularly on the adaxial part of the gutter-shaped leaf (Figure 2B-C). Water pores (Figure 2C) differ from stomata found on the leaf blade (Figure 2D-E) because they were deeply embedded below the epidermal surface, lacked or had reduced wax on their subsidiary and guard cell surfaces, and remained wide open. Moreover, water pores had a distinctive round shape unlike the elongated shape of stomata (Figure 2C-D).

**Figure 2.**
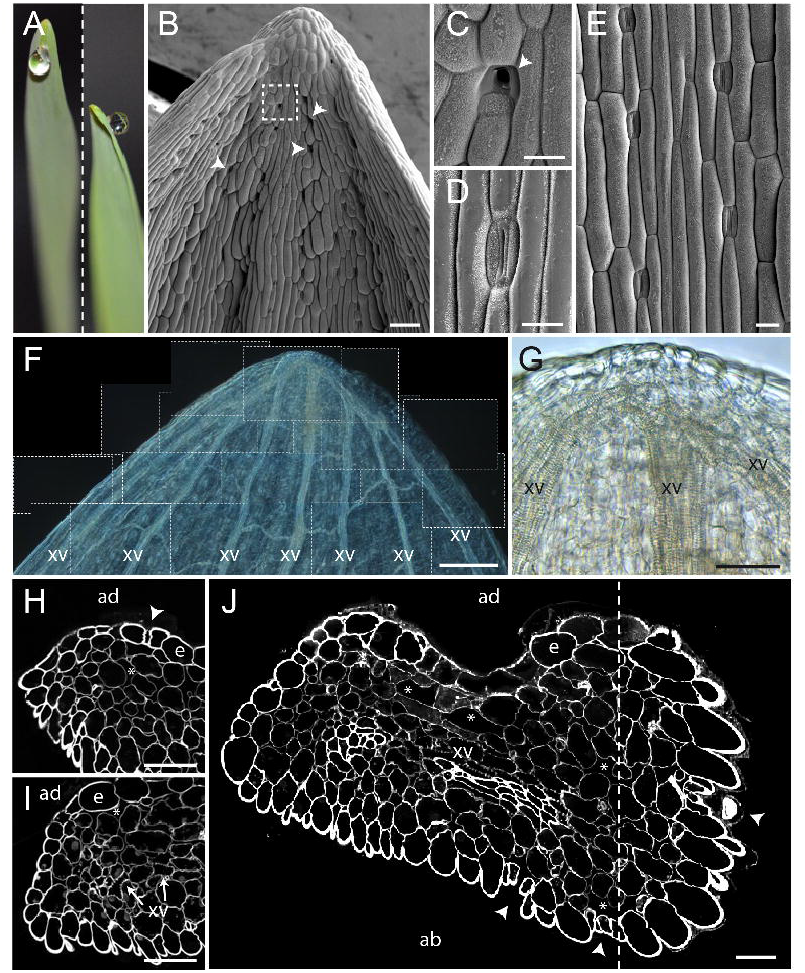
Anatomy of Barley Leaf Apical Hydathodes. **A**, Guttation droplets exuding at the leaf apex. **B-E**, Cryo-prepared leaf tip samples observed by cryo-scanning electron microscopy showing the adaxial face of the barley leaf, including a water pore close-up (C; boxed in B) and stomata (D-E). Waxes accumulate on the guard and subsidiary cells of stomata but not on water pores. **F-G**, Nomarski observation of barley leaves clarified for several weeks in a chloral hydrate solution. **F**, Mosaic of 25 individual images. **H-J**, Transverse and oblique sections of barley hydathodes. Confocal images of 1 µm-thick sections at ∼30-50 µm (H), ∼50-100 µm (I), and ∼100-150 µm (J) below the leaf apex. ad; adaxial face, ab; abaxial face of the leaf blade, e; epidermis, xv; xylem vessels. Arrowheads and asterisks indicate hydathode water pores and large spaces between epithem cells, respectively. The dashed lines indicate that this image is a mosaic of micrographs. Scale bars: 100 µm (B), 30 µm (C-E), 250 µm (F) and 50 µm (G-J).

To observe the internal tissues of hydathodes, we examined cleared leaf samples using Nomarski microscopy to enhance contrast. The leaf tip is characterized by numerous veins that converge towards the apex, connected by smaller transverse veins (Figure 2F). At approximately 30-50 µm from the tip, multiple vascular xylem endings can be found (Figure 2G). Additionally, we obtained thin transverse sections of resin-embedded material (Figure 2H-J). At 30 µm from the tip, water pores and spacious chambers are visible (Figure 2H). In sections made at 50-100 µm from the tip, the xylem vessels become visible (Figure 2I-J). Water pores are present on both adaxial and abaxial sides of the tip and open into large cavities. The xylem vessels are surrounded by a loosely arranged parenchyma with meatuses and extensive intercellular spaces referred to as the epithem. Thus, the apical hydathodes in barley exhibit the typical anatomy of passive, or so-called epithemal, hydathodes (Cerutti et al. 2019).

### *Xtt* mostly accumulates on the hydathode surface

To visualize bacterial accumulation on the leaf surface, barley leaves were dip-inoculated with *Xtt* strain UPB886 pNEO-GFP and observed via cryo-scanning electron microscopy (SEM) two days after inoculation (Figure 3). *Xtt* bacteria were visible on the leaf tip surface (Figure 3A) and detected near and on most water pores, including within their aperture (Figure 3B-C). Bacterial cells were occasionally found near stomata and substomatal cavities (Figure 3E). This indicates that the *Xtt* strain has a higher propensity to accumulate near water pores and within their aperture, which could increase the likelihood of *Xtt* infecting this organ.

**Figure 3.**
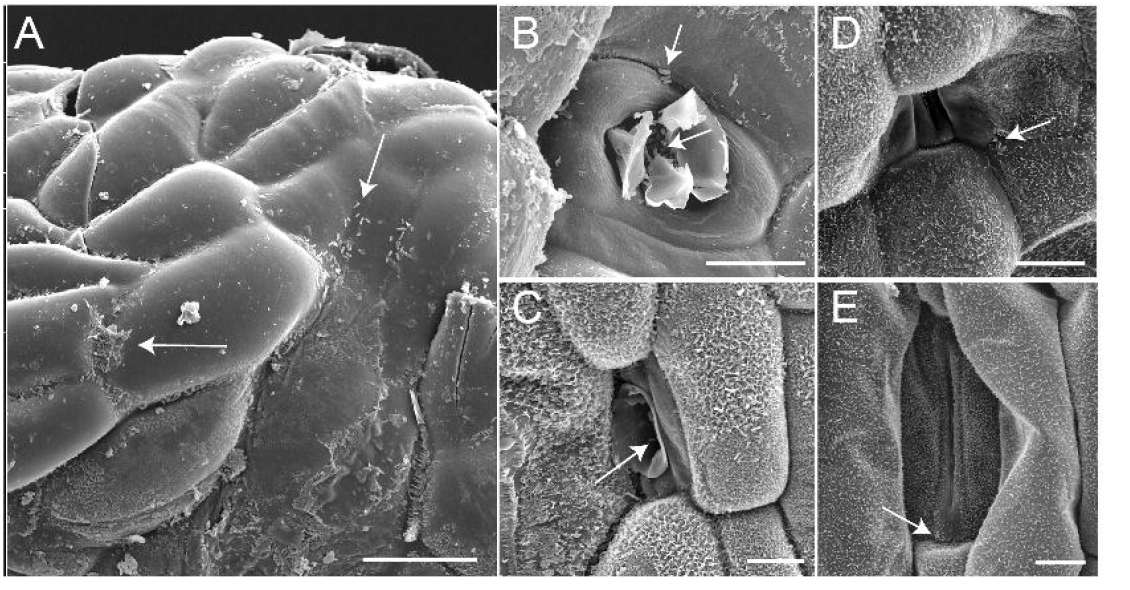
Scanning electron microscopy observations of *Xtt* strain UPB886 pNEO-GFP (A-E) at the surface of barley leaf tips two days post-dip inoculation. **A**, Bacterial cells at hydathode surface. **B-C**, Water pores with bacteria in the opening. **D**, Water pores without visible bacteria in the opening. **E**, Stomata close-up images. Waxes accumulate on both stomata guard cell and subsidiary cell surfaces but not on water pores. White arrows indicate bacterial cells. Scale bars: 25 µm (A, F), 10 µm (B-E).

### *Xtt* infects hydathodes and later colonizes xylem vessels

We then investigated the hydathode cytology at later stages of infection (Figure 4). At 6 dpi, *Xtt* bacteria were present in the extracellular spaces of the epithem (Figure 4A-B), including some epithem cells that had undergone plasmolysis (Figure 4A). However, at this stage, xylem vessels were not yet colonized. Confocal imaging of fresh leaf tips at 8 dpi revealed that GFP-tagged bacteria were primarily concentrated in the intercellular spaces of the epithem located directly beneath the water pores (Figure 4C-D). The intense blue signal detected in the vacuoles of epithem cells (between 410 and 500 nm) suggests that phenolic compounds may be present and associated with bacterial proliferation. By 13 dpi, some of the epithem cells had collapsed, and numerous xylem vessels were filled with a vivid pink fuchsine-reactive bacterial matrix (Figure 4E-F). Farther from the tip of the leaf (∼1 cm), bacteria were mainly confined to the xylem vessels (Figure 4G-H) until they could no longer be found (Figure 4I). These findings suggest that *Xtt* can effectively infect barley hydathodes and spread systemically via the xylem vessels, supporting the vascular nature of this pathogen.

**Figure 4.**
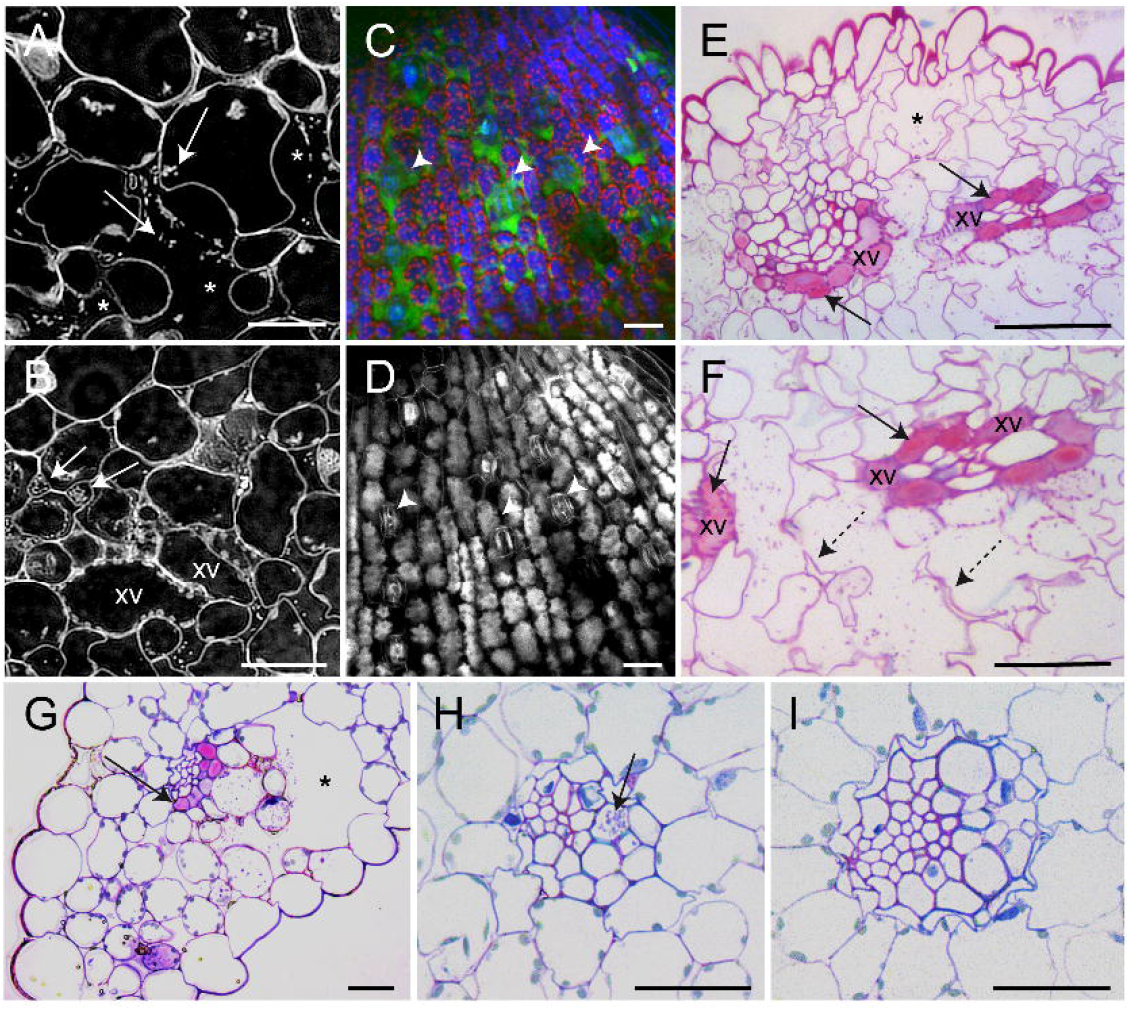
Infection of barley leaf tips by dip inoculation of *Xtt* strain UPB886 pNEO-GFP. **A-B**, Confocal micrographs of transversal 1-µm semi-thin sections of hydathodes at 6 dpi. Bacteria were mainly localized in the large intercellular spaces and in the meatuses. Bacteria were not detected in the xylem vessels (xv) at this stage. **C-D**, Confocal micrographs of fresh hydathodes at 8 dpi taken at the epithemal (C) or epidermal level (D). **D**, GFP-tagged bacteria (in green) colonized the large chambers below the water pores. Chlorophyll autofluorescence appears in red. Note the blue fluorescence from the epithem cells. Arrowheads indicate the position of water pores. **E-I** Bright-field micrographs of transversal 1-µm semi-thin sections of resin-embedded tissues at 13 dpi. Pink bacterial matrix accumulates to high levels in xylem vessels at the hydathode (E-F) and more distal parts of the leaf (G-H). **I**, Further away from the hydathodes, bacterial cells cannot be found anymore. Toluidine blue, methylene blue, and fuchsine stained plant cell walls and bacteria in pink. Dashed arrows highlight tissue degradation. Asterisks and arrows indicate large intercellular spaces and accumulation of bacterial cells and/or matrix, respectively. Scale bars: 20µm.

## Discussion

In this study, we examined the ability of *X. translucens* pv. *translucens (Xtt)* to colonize barley hydathodes. We developed a protocol that closely parallels natural barley infection by *Xtt*. We established the detailed anatomy of apical hydathodes in barley leaves and provided a microscopic description of leaf infection by *Xtt*.

### Typical epithemal hydathodes are present on barley leaves

Barley hydathodes present on leaf tips mediate abundant guttation in lab conditions. The reduced waxes on the epidermis and the cup shape may contribute to the stabilization of guttation droplets at the apex(Cerutti et al. 2019). These hydathodes exhibit all the typical features of epithemal hydathodes, including water pores, epithem, and an interconnected vascular network (Cerutti et al. 2019; Dieffenbach et al. 1980; Jauneau et al. 2020; Perrin 1972). Water pores exhibit several distinct morphological features compared to stomata, including greater aperture, irregular distribution on the leaf surface, and the absence or low accumulation of cuticular waxes at the pore and neighboring cell wall surfaces. The waterpores are also positioned deep below the epidermal surface and give access to a lacunar epithem. Previous studies reported that wall vessels within the hydathodes are not lignified and exhibit some porosity and that the vessel walls were thin near the numerous pits (Dieffenbach et al. 1980). Holes were also detected in the pit membrane of xylem vessels in other monocots such as *Brachypodium* and sugarcane (Jauneau et al. 2020). These characteristics of epithem and vessels should facilitate the propagation of adapted bacterial pathogens from the hydathode to the xylem.

### *Xtt* is one of the few demonstrated hydathode pathogen

In this study, we established the inoculation procedure to study the infection of barley hydathodes by *Xtt* under controlled conditions. We have shown that this strain can successfully invade and proliferate within hydathodes. The distinctive observation of BB symptoms in lab conditions using a non-invasive protocol reflects the capacity of *Xtt* to transition from the hydathodes to the vascular tissues and cause systemic disease symptoms as it likely occurs in the field. Such non-invasive protocols such as dip or spray inoculations are valuable because they do not bypass the plant’s natural anatomical and immune barriers (Cerutti et al. 2017; Nino□Liu et al. 2005; Whalen et al. 1991) and because they facilitate the study of the complete life cycle of the pathogen. Bacterial entry through water pores may be passive and dependent on the guttation cycles, which are driven by the balance between root pressure and leaf evapotranspiration (Cerutti et al. 2019). In support of this hypothesis, *Xcc* motility mutants are not affected in their capacity to colonize cauliflower hydathodes (Luneau et al. 2022b). Once inside hydathodes, bacteria face various immune responses, as best described in the *Xcc*/Brassicaceae pathosystem: the Hrp type III secretion system and its effector proteins dampen immunity in cauliflower (Cerutti et al. 2017). Moreover, several pathogen-associated molecular pattern (PAMP) receptors, associated signaling components and defense-related genes are expressed in Arabidopsis hydathodes (Beck et al. 2014; Paauw et al. 2023; Routaboul et al. 2024). *ZAR1*-, *xopJ6*- and *SUT1*-dependent effector-triggered immunity was also recently demonstrated to be effective inside hydathodes to limit *Xcc* proliferation (Lauber et al. 2024; Paauw et al. 2023; Taks et al. 2025). As for barley, pathogenesis-related proteins have been found in the guttation fluid of healthy barley leaves, indicating some basal level of defenses at the hydathode level (Grunwald et al. 2003). Thus, to date, the immunity faced by *Xtt* in hydathodes and the vasculature remains to be determined and characterized in order to better understand and control this important crop disease.

## Acknowledgements

We wish to thank Bruno Payre from the CMEAB facility of Toulouse University for the preparation of the cryosamples.

## Funding

The authors are grateful to the funding sources: JMJ was supported from a COST SUSTAIN STSM award (FA1208-060317-082568), a Fulbright Scholar Award from the US Fulbright Commission to Belgium/Luxembourg, a USDA NIFA Postdoctoral Fellowship (#2016-04706), USDA NIFA award no. 2018-67013-28490 through the NSF/NIFA Plant Biotic Interactions Program and the Ethel and O.N. Endowment at UW–Madison. AC, AJ and LDN are supported by the NEPHRON ANR project (ANR-18-CE20-0020-01). A.C. was funded by a PhD grant from the French Ministry of National Education and Research. This study is set within the framework of the ‘Laboratoires d’Excellences’ (LABEX) TULIP (ANR-10-LABX-41) and of the ‘Ecole Universitaire de Recherche’ (EUR) TULIP-GS (ANR-18-EURE-0019). Authors benefited from the COST action CA16107 EuroXanth.

## Author contributions

AJ, CB, JMJ and LDN conceived and designed the experiment; AC established the hydathode infection protocol; AC, JB and JMR performed the inoculations; AJ, AC, JB and MCA performed the microscopy; AJ and LDN assembled the figures; AJ, JB, JMR, JMJ and LDN wrote the manuscript. All authors contributed to the final version.

## Notes

### Competing Interest Statement

The authors have declared no competing interest.

